# Deep Mendelian Randomization: Investigating the Causal Knowledge of Genomic Deep Learning Models

**DOI:** 10.1101/2022.02.01.478608

**Authors:** Stephen Malina, Daniel Cizin, David A. Knowles

## Abstract

Multi-task deep learning (DL) models can accurately predict diverse genomic marks from sequence, but whether these models learn the causal relationships between genomic marks is unknown. Here, we describe Deep Mendelian Randomization (`DeepMR`), a method for estimating causal relationships between genomic marks learned by genomic DL models. By combining Mendelian Randomization with *in silico* mutagenesis, `DeepMR` obtains local (locus specific) and global estimates of (an assumed) linear causal relationship between marks. In a simulation designed to test recovery of pairwise causal relations between transcription factors (TFs), `DeepMR` gives accurate and unbiased estimates of the ‘true’ global causal effect, but its coverage decays in the presence of sequence-dependent confounding. We then apply `DeepMR` to examine the global relationships learned by a state-of-the-art DL model, BPNet [Avsec et al., 2020], between TFs involved in reprogramming. `DeepMR`’s causal effect estimates validate previously hypothesized relationships between TFs and suggest new relationships for future investigation.

## 1 Introduction

Deep learning (DL) has achieved success predicting genomic marks^1^ such as transcription factor (TF) binding [Alipanahi et al., 2015, Zhou and Troyanskaya, 2015], chromatin accessibility [Zhou and Troyanskaya, 2015, Kelley et al., 2016], histone marks [Yin et al., 2019], RNA binding protein (RBP) binding [Alipanahi et al., 2015, Pan and Shen, 2017, Gandhi et al., 2018, Zheng et al., 2018] and splicing [Jaganathan et al., 2019, Cheng et al., 2021] from DNA (or RNA) sequence. These models typically achieve high predictive accuracy and recognize sequence features that match those found by orthogonal experiments such as SELEX [Tuerk and Gold, 1990]. In particular, multi-task models such as DeepSEA [Zhou and Troyanskaya, 2015] and BPNet [Avsec et al., 2020] can accurately predict multiple genomic marks simultaneously. Here we ask: do such multi-task models, through learning to predict multiple marks jointly, gain an implicit understanding of mechanistic, causal relationships between marks?

We attempt to answer this question by developing Deep Mendelian Randomization (`DeepMR`). `DeepMR` combines *in silico* mutagenesis with Mendelian randomization [Lawlor et al., 2008], an instrumental variable approach for causal inference, to estimate learned causal effects in genomic DL models. `DeepMR` obtains local (sequence level) and global (genome level) estimates of (an assumed) linear causal relationship between pairs of marks learned by a multi-task genomic prediction model. `DeepMR` draws on four threads of work spanning machine learning and statistical genetics.

### DL for Functional Genomics

A major objective in functional genomics is mapping sequence-to-function relationships between genotype and molecular phenotypes, typically leveraging large-scale observational data from high-throughput assays such as ChIP-seq [Johnson et al., 2007, Barski et al., 2007, Robertson et al., 2007, Mikkelsen et al., 2007], DNase-seq [Song and Crawford, 2010], and ATAC-seq [Buenrostro et al., 2015]. Understanding this mapping enables 1) better understanding of epigenomic regulation, 2) variant interpretation, and 3) more accurate prediction of downstream traits. However, achieving these goal requires decoding complex relationships between high-dimensional genomic sequence inputs and interrelated outputs from large, noisy datasets. Encouraged by DL models’ ability to overcome similar challenges in the fields of computer vision and natural language processing, genomics researchers have trained DL models on functional genomics datasets with substantial success.

Early work showed that DL could predict sequence-to-function relationships accurately and demonstrated their promise for identifying trait-associated variants. DeepBind [Alipanahi et al., 2015], one of the earliest DL sequence-to-function classifiers, outperformed then state-of-the-art models at predicting TF binding and RBP binding from sequence. DeepBind and other classification models—e.g. DeepSEA [Zhou and Troyanskaya, 2015] and Basset [Kelley et al., 2016]—also identified trait-associated variants with higher accuracy than previous methods. More recent work has leveraged DL models to improve our understanding of epigenomic regulatory logic. In particular, Avsec et al. [2020] trained a regression model, BPNet, to predict the binding of four TFs and used it to dissect the motif-based regulatory grammar that governs their binding. Together, these papers illustrate the promise of DL models for not only predicting function from sequence but also improving our understanding of epigenomic regulation and ability to anticipate disease risk. In our work, we seek evidence that genomic DL models learn meaningful high-level relationships between output marks.

### Model Interpretability

Local interpretation methods characterize how specific input (sequence) features influence predictions or intermediate layer activations (e.g., saliency maps [Simonyan et al., 2013], guided back-propagation [Springenberg et al., 2014], DeepLIFT [Shrikumar et al., 2017], and DeepSHAP [Lundberg and Lee, 2017]). Even DeepLIFT, which was designed with genomic DL in mind, focuses on interpreting individual model predictions for a single output rather than discovering relationships between outputs and is therefore complementary to our work.

Closer to our work, Koo et al. [2021]’s Global Importance Analysis (GIA) assesses the global effect size of different patterns on model predictions. While resembling `DeepMR` in terms of its focus on global effects, GIA allows users to test narrower hypotheses about specific features such as motifs and uses synthetic instead of observed sequences. As such, GIA is also complementary to `DeepMR`, potentially providing a method for uncovering specific patterns that explain higher level relationships discovered by `DeepMR`.

Saturation *in silico* mutagenesis characterizes how a model’s predictions for an input change as a result of all possible point mutations to the input. Saturation mutagenesis has been used to assess the learned representations of genomic DL models such as DeepBind [Alipanahi et al., 2015], cDeepBind [Gandhi et al., 2018], DeepSEA [Zhou and Troyanskaya, 2015], and Basset [Kelley et al., 2016]. Here, we use saturation mutagenesis (with uncertainty estimates generated using Deep Ensembles [Lakshminarayanan et al., 2016]) to generate a set of estimated variant *effect sizes* which we then provide as input to Mendelian randomization.

### Uncertainty Estimates and Coverage of DL Predictions

Many methods for obtaining uncertainty estimates from DL models exist [Abdar et al., 2021]. Our work is not focused on testing different uncertainty estimation methods so we chose Deep Ensembles [Lakshminarayanan et al., 2016], which, despite their simplicity, consistently perform well in empirical comparisons [Lakshminarayanan et al., 2016, Hirschfeld et al., 2020]. Despite this, Kuleshov et al. [2018] found that uncertainty estimates from Deep Ensembles were often miscalibrated but could be rescued using isotonic regression (a solution we adopt here).

### Mendelian Randomization

Mendelian randomization (MR) is a technique for estimating linear causal effects in the presence of potential unobserved confounders. While MR is typically used to estimate inter-phenotype causal effects from population-scale observational data (i.e., genome-wide association studies, GWAS), here we explore its application to estimating causal effects implied by model-generated “data” but with *in silico* mutations taking the place of true genetic variants.

MR only produces valid causal effect estimates under specific assumptions (Figure 1 under <u>Estimate</u>) Lawlor et al. [2008]. Let *Z* be a variable we intend to use as an instrument (a genetic variant for example), *X* a purported cause (*exposure*), and *Y* a purported effect (*outcome*), and suppose that there may be unobserved confounding between *X* and *Y*, denoted by *U*. Then, MR gives an unbiased estimate of the causal effect of *X* on *Y* if:

1. *Z* is independent of *U* (**Unconfoundedness**),
2. *Z* is not independent of *X*, and
3. *Z* only influences *Y* through *X* (**Exclusion Restriction**).

**Figure 1:**
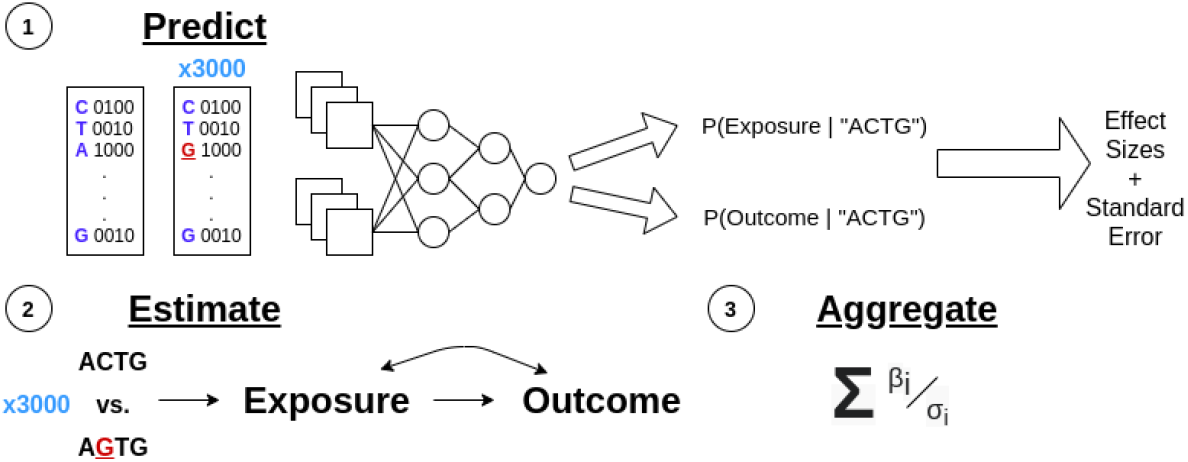
Graphical representation of Deep MR’s high-level steps combining *in silico* mutagenesis and Mendelian randomization (see Section 2.1). <u>Predict</u> corresponds to steps 1 through 4. <u>Estimate</u> corresponds to step 5. <u>Aggregate</u> corresponds to step 6

Recently developed MR methods such as Robust Adjusted Profile Score [Zhao et al., 2018], MR-Egger [Bowden et al., 2015], and the modal-based estimator [Burgess et al., 2018] leverage multiple instruments to relax some of these assumptions without compromising the validity of results. In this work, we estimate causal effects using a robust variant of MR-Egger with the goal of being robust to invalid instruments. MR-Egger seeks robustness to violations of Exclusion Restriction, otherwise known as horizontal pleiotropy in statistical genetics [Hemani et al., 2018].

## 2 Methods

### 2.1 Algorithm Overview

`DeepMR` estimates causal effects between variables predicted by a multi-task model. It takes a trained, calibrated (regression or classification) model that outputs predictive means and standard errors and a set of one-hot encoded sequences as input. It outputs local, sequence-specific causal effects and global, exposure/outcome-specific causal effects. It accomplishes this (see Figure 1 for a visual depiction) via the following steps for each exposure/outcome pair:

1. Randomly sample sequences to predict exposure and outcome values for “reference sequences”.
2. Perform saturation *in-silico* mutagenesis for each reference sequence to generate (sequence length × alphabet size − 1) mutated sequences per reference sequence.
3. For each set of pairs of mutant and reference sequences, generate predictive means and standard errors for exposure and outcome features.
4. Generate (sequence length×alphabet size−1) *effect sizes* by subtracting each reference sequence’s predictive mean from the corresponding mutated sequences’ predictive means. Additionally, compute the standard errors of these differences.
5. Filter instruments by effect size based on a *z*-score threshold (Assumption 2) to only include those that are strongly associated with the exposure.
6. Estimate a per-exposure, per-sequence region causal effect by running MR on the remaining effect sizes and their standard errors.
7. Estimate overall per-exposure causal effects using a random effects meta-analysis across sequence regions (loci).

### 2.2 Simulation

Our simulation is inspired by Finkelstein et al. [2020] but tailored to test `DeepMR`’s ability to estimate the strength of the causal relationship between exposure and outcome TFs when binding to simulated *L* = 100bp DNA sequences. The exposure TF’s binding affinity, *c_e_*, is determined primarily by the probability of the TF (represented as a position weight matrix, PWM) binding anywhere on the sequence (see Appendix 5), *p_e_*,

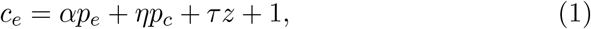

where *p_c_* is the binding probability of an optional confounder TF, and *z* ~ Bernoulli(0.5) is an optional sequence independent confounder. By contrast, the outcome TF’s binding affinity *c_o_* is a multiplicative function of both the strength of its own motif match and the strength of the exposure’s, i.e.

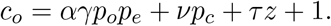

Here the effect size *γ* represents the influence of the exposure’s binding on the outcome’s binding in raw counts space. *γ* is not the *true causal effect* because the true CE is defined in Anscombe-transformed rather than raw counts space. *γ* is sampled (once per simulation run) from an equal proportion mixture of two normals with means 10 and 1 (and variance 0.5), in order to test `DeepMR`ś ability to differentiate between two clusters of CEs, one much lower than the other. We sample α from *N*(100, 3) (once per simulation run) and fix *η* = 20, *ν* = 30 and *τ* = 25.

The simulation model corresponds to a causal effect of the exposure TF on the outcome TF: with no exposure TF binding there can be no outcome TF binding. When present, both types of confounding influence exposure and outcome counts multiplicatively.

Finally to represent experimental noise, counts are Poisson distributed with mean equal to the affinity values. We did not use a negative binomial since we expect the random sequence generation process will naturally induce overdispersion. Finally, we Anscombe transform the raw counts [Finkelstein et al., 2020].

Length 100 sequences are sampled uniformly at random. For each TF, with 50% probability we insert a subsequence sampled from its PWM at a random position. To assign a binding probability we convolve the TF’s PWM over the sequence and apply the soft-or function (see Appendix 5).

We considered four different scenarios: 1) no unobserved confounding, 2) sequence-based unobserved confounding, 3) non-sequence-based unobserved confounding, and 4) both types of confounding in tandem. Sequence-dependent unobserved confounding adds an additional TF (and corresponding) motif which influences the binding strength of both exposure and outcome TFs.

We train ensembles of convolutional neural network (CNN) models on the data produced in each scenario and use them, combined with held-out test sets, as inputs for `DeepMR`.

#### True Causal Effect Computation

To assess the quality of our method, we need to compare its estimates to the ground truth. `DeepMR` estimates the effect of a unit change in the exposure on the outcome by using single point mutations that meaningfully affect the exposure as instruments. Our simulation can provide us with the true affinity for any given mutated sequence, which we leverage to compute true sequence-region level causal effects. For a given sequence which contains the exposure motif, the true causal effect is found by regressing the effect of all point mutations to bases within the exposure motif on the outcome on the corresponding effects on the exposure. This is similar to the two-stage least squares MR method [Angrist and Imbens, 1995] where all mutations within the the exposure motif are assumed to be valid instruments.

#### Simulation & Model Parameters

In all simulation runs, we used PWMs representing motifs for the GATA (exposure), TAL1 (outcome), and SOX2 (confounder) transcription factors, all drawn from ENCODE’s motif database [Kheradpour and Kellis, 2014] and sampled using the simdna library (https://github.com/kundajelab/simdna).

To model this data, we trained 3-layer convolutional neural network [LeCun et al., 1995] with 15 filters per convolutional layer and a filter width of 7 for maximum 100 epochs with early-stopping. The three convolutional layers were followed by 2 hidden layers of width 30. Models were trained using an MSE loss to predict the Anscombe-transformed counts for the exposure and outcome jointly.

Code for all experiments can be found at https://github.com/an1lam/deepmr.

## 3 Results

We first assess `DeepMR` on simulated data where we know the ground-truth relationship between the modeled TFs. We then apply `DeepMR` to determining the causal relationships between four TFs involved in pluripotency.

### 3.1 Simulation

#### DeepMR accurately estimates global CEs in all cases

We evaluated `DeepMR`’s local and global CE estimates in the one unconfounded and three confounded scenarios (see Methods). In each scenario, we performed causal effect estimation (including learning the sequence-to-binding CNN ensemble) for 50 simulations using 10000 training sequences and 1000 test sequences for CE estimation.

Our CNN models achieved *R*^2^ validation accuracy averaging around 0.8 for (transformed) exposure counts and 0.7 for (transformed) outcome counts. For the causal inference we assessed two metrics: accuracy of global CEs and coverage of local CE 95% confidence intervals. We judged accuracy of global CEs in terms of the correlation between the global causal effect estimates and the average of the true global causal effects and the frequency at which the CE estimate ±2*τ* capture said average across 50 simulations. `DeepMR` accurately estimates true global CEs in all cases (Fig. 2, Table 1 for *R*^2^ accuracy values). In the unconfounded and non-sequence confounding cases, we see near-perfect agreement between estimated and true global CEs. In the sequence-dependent confounding case, `DeepMR` more often underestimates true CEs, although usually by less than ones SE, suggesting that the influence of the unlabeled SOX motif score on the exposure and outcome label values biases `DeepMR`’s global CE estimates towards 0.

**Figure 2:**
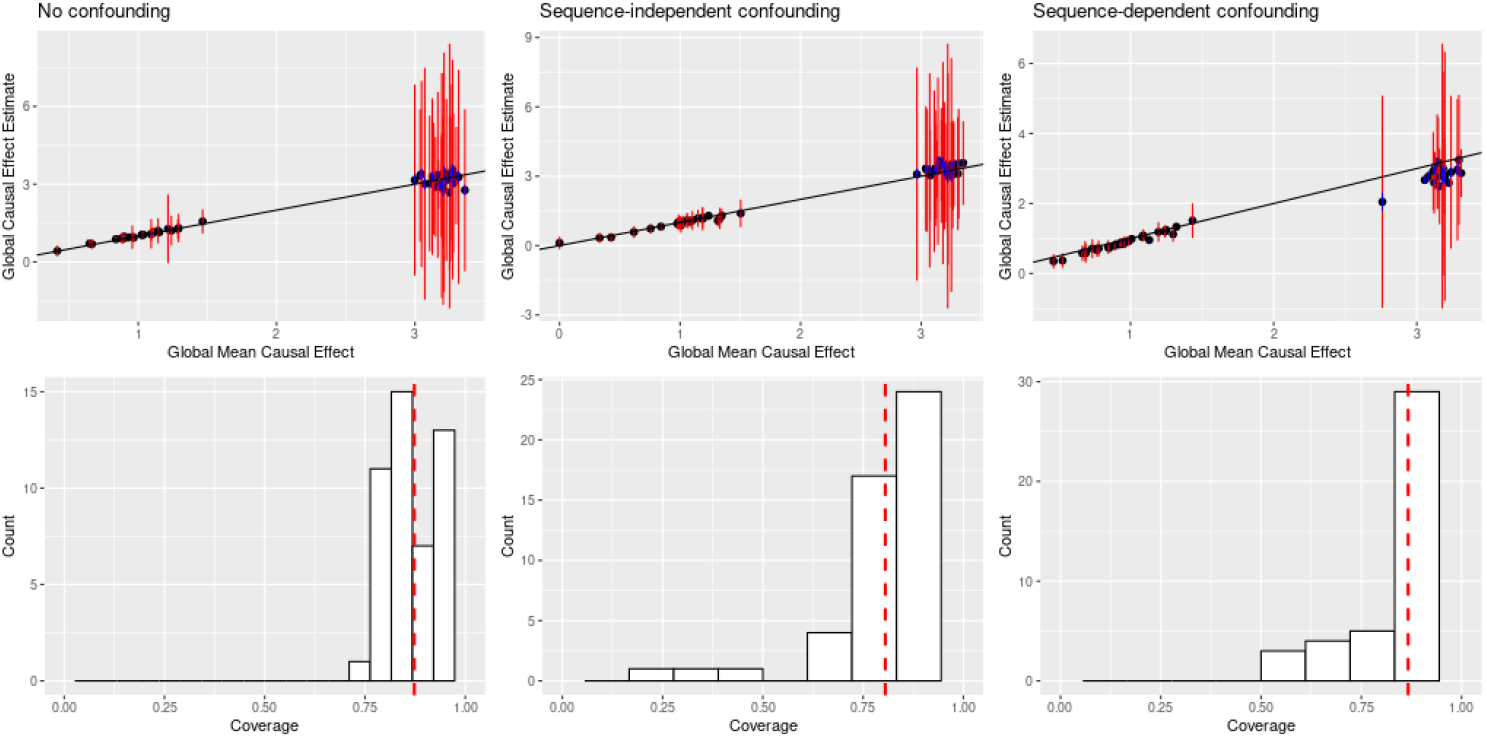
In simulations, `DeepMR` estimates causal effects between TFs even in the presence of unobserved confounding. **Top row:** true vs. estimated global causal effects (CEs) across 50 rounds for unconfounded, random confounded, and sequence-dependent confounded cases respectively. Blue bars denote ±2*σ* where *σ* denotes the standard error of the mean and orange bars denote ±2*τ* where *τ* denotes the between-region standard deviation. **Bottom row:** local CE coverage (how often the true CE is in the 95% confidence interval) across the three experiments (same order) with the red line denoting average coverage.

**Table 1:** `DeepMR` estimates causal effects (CE) accurately with high coverage. Accuracy is Pearson *R*^2^. Local corresponds to CEs for individual regions, global for the meta-analysis mean.

|  | Global CE Accuracy | Global CE Coverage | Local CE CI Coverage |
| --- | --- | --- | --- |
| Unconfounded | 0.97 | 0.90 | 0.87 |
| Random | 0.98 | 0.96 | 0.81 |
| Sequence | 0.98 | 0.68 | 0.87 |
| Both | 0.78 | 0.92 | 0.81 |

#### DeepMR’s coverage decays in the presence of sequence-dependent confounding

We judged coverage of global CEs by measuring the fraction of ±2*τ* intervals that capture the true global CE. We judged coverage of local CEs by examining the distribution of 95% confidence interval coverage across 50 simulations in the four scenarios. `DeepMR` performs better in the unconfounded and random confounding scenarios. While average coverage (see Supplementary Table 1) is relatively constant across scenarios, in Fig. 2 we observe a longer tail of low coverage values in the random and sequence-dependent confounding scenarios. Furthermore, global CE coverage in the sequence-dependent and scenario is much lower. Together, these results suggest that `DeepMR` can somewhat underestimate variance in confounded scenarios and produces more calibrated local CE estimates in cases where there is minimal or no sequence-dependent confounding.

### 3.2 Estimating causal effects between four TFs involved in reprogramming

Given the promising results on simulated data, we applied `DeepMR` to detecting CEs between four TFs involved in induced pluripotent stem cell (iPSC) reprogramming: Oct4, Sox2, Nanog, and Klf4. We used the ChIP-nexus data and model (BPNet) previously described in [Avsec et al., 2020] but trained a 5-component ensemble. We closely followed the data processing and model training process used in the original paper, described in full at the BPNet repository (https://github.com/kundajelab/bpnet). We calibrated the resulting Deep Ensemble with isotonic regression using validation data. We computed local CE estimates for all TF pairs on 2000 randomly sampled sequences in the validation set. These estimates were used to compute global estimates for each TF pair via meta-analysis.

#### DeepMR validates previously hypothesized and suggests new relationships between TFs

Based on an orthogonal approach (TF cooperativity analysis), Avsec et al. [2020] postulate a positive directional effect of a composite Oct4-Sox2 binding motif on the binding of Nanog and Klf4. As a test of `DeepMR`’s ability to discover such relationships while making fewer assumptions about their functional form, we sought to replicate this finding. While the BPNet approach does not produce quantitative overall estimates of directional effects, it enabled them to make two hypotheses about directionality (see Extended Data Fig. 6). These were 1) Sox2 and Oct4 act on Nanog and 2) that Oct4 and Sox2 act on each other via a composite motif. To replicate these findings, we computed the global CEs for all 12 pairs of TFs. We largely recapitulate Avsec et al. [2020] (Figure 3), finding that Sox2 and Oct4 both have a strong positive estimated CEs on each other’s binding and on the binding of Nanog. In the latter case, the 2*τ* range does include 0, suggesting high variability across loci. In general, we observe high variability across sequence regions, reflected by the generally large ±2*τ* ranges. This also matches Avsec et al. [2020]’s observation that effects vary across sequence space and in particular with different motif spacings.

**Figure 3:**
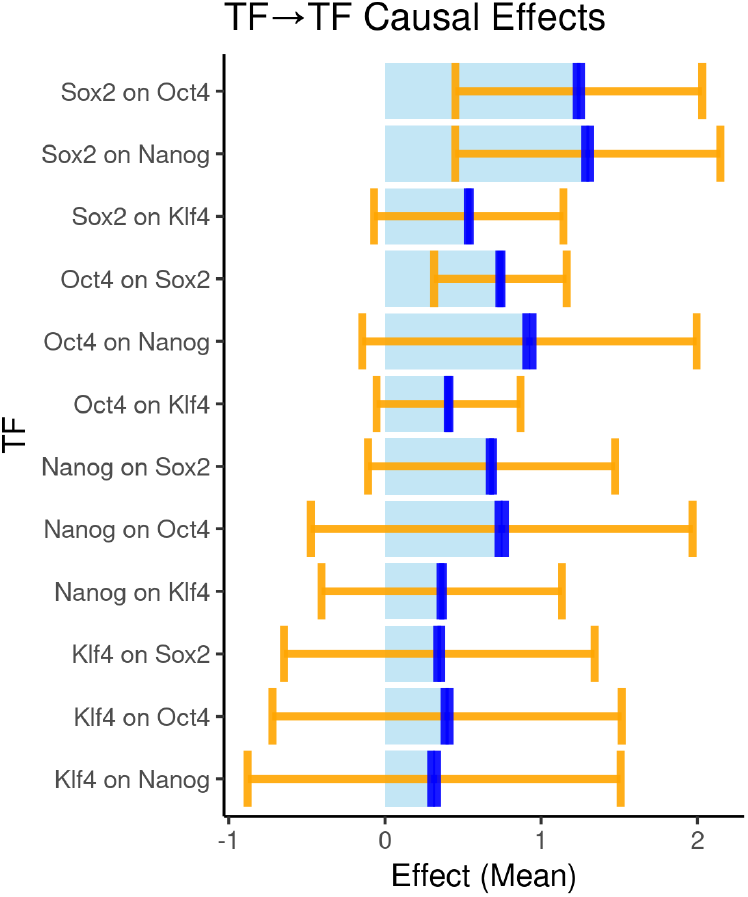
Global CEs for all pairs of TFs predicted by BPNet with ±2τ (orange) and ±2σ (blue) ranges around the mean estimate.

`DeepMR` suggests additional hypotheses that could be validated by in future experimental work. As one example, `DeepMR` predicts that Sox2 acts on Klf4 more strongly than the reverse.

## 4 Discussion

`DeepMR` estimates the magnitude of causal relationships between outputs of multi-task genomic DL models in order to hypothesize specific models of gene regulation. `DeepMR` can recover CEs in the presence of unobserved confounding in simulation and validates purported and identifies new putative relationships between four important TFs involved in reprogramming. While `DeepMR` shows promise, it does have several known limitations.

### Resource Requirements

Since `DeepMR` relies on *in silico* mutagenesis, generating the data for estimating global CEs is computationally intensive, taking approximately one day to run for the BPNet hold-out set in our experiments. One could incorporate speed-ups such as those of Nair et al. [2020] or leverage attribution tools such as saliency maps, DeepLIFT [Shrikumar et al., 2017] or DeepSHAP [Lundberg and Lee, 2017] that can efficiently approximate *in silico* mutagenesis.

### Model Calibration

MR Egger requires properly calibrated effect size and standard error estimates for each instrument. Our ensemble-based approach to uncertainty estimation tends to produce somewhat over-confident estimates as measured by the metrics proposed by Kuleshov et al. [2018]. We apply and recommend isotonic regression [Barlow and Brunk, 1972] to remedy this.

### Violation of MR assumptions

For MR to return unbiased causal effect estimates, the underlying data-generating process and our model’s proxy for it must both adhere to the three MR assumptions (1) and there must be an at least approximately linear relationship between exposure and outcome. In the statistical genetics setting, these assumptions can be justified in part by claims about the relationship between genotype, which is determined prenatally, and potential confounders and phenotypes, which tend to manifest post-natally, assuming population structure is accounted for. We cannot fall back on these justifications for sequence-to-function relationships. Instead, we must re-examine each of these assumptions to determine whether they can be expected to hold. Assumption 2 is easily satisfied because by filtering instruments based on their relationship to the exposure (see Section 2.1), whereas the unconfoundedness (Assumption 1), exclusion restriction (Assumption 3), and linearity assumptions have the potential to be violated.

Under classical MR assumptions, estimates will only be unbiased if all instruments are independent of unobserved confounders. Potential unobserved confounders fall into two categories: sequence-dependent and sequence-independent. Classical MR (i.e. inverse-variance weighting) should control for sequence-independent confounding. Potential sequence-dependent confounders include other TFs, chromatin features or an uncorrected assay bias such as GC-bias. Such confounders additionally violate the exclusion restriction assumption by providing a causal pathway from instrument (mutation) to outcome not mediated by the exposure TF. However, our use of MR Egger provides some additional robustness to such violations so long as the InSIDE assumption holds (see 1). Indeed, our simulation experiments (section 3.1) showed remarkable robustness to the effects of both types of confounding.

MR correctly estimates causal effects when all relationships – instrument to exposure and exposure to outcome – are linear, which may not be the case. For example, given strong TF binding cooperativity, knocking out one TF’s binding will knock out the other’s entirely, violating linearity. Fortunately, we only require the weaker condition of local linearity. Each of our effect sizes is derived from a single mutation, so `DeepMR` behaves correctly so long as the relationships stay linear within a local neighborhood.

In summary, `DeepMR` relies on specific assumptions about model quality and the true causal relationships. The former can be expected to increase as genomic datasets grow. The latter suggests relaxing some of these assumptions via more advanced MR methods or developing tools to detect when assumptions are violated.

In the future, we will aim to combine `DeepMR` with a causal network inference method such as our recent bimmer model [Brown and Knowles, 2020] to explicitly account for the influence of other assayed TFs on each pair. `DeepMR` would also benefit from accompanying tools for diagnosing when model-generated data deviates from or violates MR assumptions.

## 5 Acknowledgments

We thank Brielin Brown, Megan Schertzer, Laura Pereira, Udai Nagpal, Collin Wang, and Nasrine Metic for discussion of the idea, experiments, and analysis. We thank Milan Cvitkovic and Pablo Cordero for feedback on the manuscript.

## Appendix A Computing Binding Probabilities

Given a PWM and a sequence, denoted by *s* ∈ {0, 1}^4×*L*^, we want to obtain the probability of observing binding at any of the sequence’s *M*-length subregions under the assumption that binding probabilities are independent across the subregions. We first compute the probability of binding at each subregion follow the process described by Finkelstein et al. [2020], a modified version of the energy-based model of binding proposed in Zhao et al. [2012].

To aggregate probabilities across positions into a single sequence binding probability, we let *p_i:i+M_* denote the probability of the TF binding at sequence subregion spanning nucleotide *i* to *i* + *M* and compute the probability of binding somewhere on the sequence as

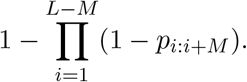

That is, the probability of the TF binding somewhere is one minus the probability of it not binding anywhere. This operation is sometimes known as “soft-or”.

## Footnotes

1 Following Trynka et al. [2013], we define a “mark” as a position in the genome where the number of reads from an epigenomic assay is significantly above background.

